# Whole-Genome Sequences of the Severe Acute Respiratory Syndrome Coronavirus-2 obtained from Romanian patients between March and June of 2020

**DOI:** 10.1101/2020.06.28.175802

**Authors:** Mihaela Lazar, Odette Popovici, Barbara Mühlemann, Tim Durfee, Razvan Stan

## Abstract

Impact of mutations on the evolution of Severe Acute Respiratory Syndrome Coronavirus-2 (SARS-CoV-2) are needed for ongoing global efforts to track and trace the current pandemic, in order to enact effective prevention and treatment options. SARS-Co-V-2 viral genomes were detected and sequenced from 18 Romanian patients suffering from coronavirus disease-2019. Viral Spike S glycoprotein sequences were used to generate model structures and assess the role of mutations on protein stability. We integrated the phylogenetic tree within the available European SARS-Co-V-2 genomic sequences. We further provide an epidemiological overview of the pre-existing conditions that are lethal in relevant Romanian patients. Non-synonymous mutations in the viral Spike glycoprotein relating to infectivity are constructed in models of protein structures. Continuing search to limit and treat SARS-CoV-2 benefit from our contribution in delineating the viral Spike glycoprotein mutations, as well as from assessment of their role on protein stability or complex formation with human receptor angiotensin-converting enzyme 2. Our results help implement and extend worldwide genomic surveillance of coronavirus disease-2019.

## Introduction

In December 2019, in China, an outbreak started due to a novel coronavirus strain that causes a severe illness, the coronavirus disease 2019 (COVID-19). The virus, SARS-CoV-2, has since spread globally [1]. Data reported to European Center for Disease Control show that clinical presentation of COVID-19 ranges from no symptoms to severe pneumonia, and that many cases have led to death [2]. According to available data, 32 % of the diagnosed COVID-19 cases in the EU/EEA have been hospitalized, and 4% thereof had severe form of illness [3]. Additionally, hospitalization rates have been significantly higher for adults 65 years and older [4]. The first positive case of infection with SARS-CoV-2 in Romania was confirmed on 26^th^ of February 2020. With the ongoing spread of the virus, it is becoming critical to detect and classify the virus in patient samples. As such, full genome characterization of this virus is instrumental for updating diagnostics criteria and assessing viral evolution. To assess its genetic variation in a South-Eastern European population, we herein generated genome sequences using metagenomic sequencing, from 18 Romanian patients. We delineated the non-synonymous mutations and constructed models of the protein structures from these sequences. We explicitly focused on the spike (S) glycoprotein because it mediates infection of human cells, and is the main target of most vaccine strategies and antibody-based therapeutics [5].

## Materials and Methods

### Collection and processing of samples

Nasopharyngeal and oropharyngeal swabs were collected from individuals from different Romanian geographical areas. Total nucleic acid extraction was performed using the Maxwell**®** RSC Viral Total Nucleic Acid Purification Kit (Promega, USA) as described by the manufacturer. SARS-CoV-2 nucleic acid was detected using the E gene assay as the first-line screening tool, followed by confirmatory testing with the RdRp gene assay [6]. The RiboZero Gold rRNA depletion protocol was used to remove human cytoplasmic and mitochondrial rRNA. The total RNA quantity and integrity were measured with the Qubit RNA Assay Kit (Invitrogen, Carlsbad, CA, USA). The TrueSeq Stranded Total RNA Library Prep Gold kit along with the IDT for Illumina TruSeq RNA UD Indexes were used for sequencing-ready library preparation. RNA fragmentation, first and second-strand cDNA synthesis, adenylation, adapter ligation and amplification were done according to the TruSeq Stranded Total RNA protocol. After amplification, the prepared libraries were quantified, pooled and loaded onto Illumina MiSeq DNA Sequencer. Sequence data from each sample was aligned to the SAR-CoV-2 reference genome (GenBank accession number: NC_045512.2) and variants called using SeqMan NGen (DNASTAR, Madison, WI, USA). Alignments were manually inspected to confirm variant calls and the viral sequence from each sample exported with SeqMan Pro (DNASTAR).

### Data mining for structures

Atomic coordinates of the template, the novel SARS-CoV-2 spike protein in complex with Receptor Binding Domain (RBD) of the human Angiotensin-converting enzyme 2 (hACE2), PDB: 6LZG, were retrieved from RCSB protein data bank. Spike protein sequences from our genomes were modeled using I-Tasser [7] and the quality of models of Spike S proteins was assessed with MolProbity [8]. Full length models were superimposed over template and root mean square deviations (RMSD) of carbon alpha backbones were determined using PyMol (PyMOL Molecular Graphics System, Version 2.0 Schrödinger, LLC).

### Mutational analyses and molecular docking

Site Directed Mutator (http://marid.bioc.cam.ac.uk/sdm2/) was used to assess the impact of non-synonymous single, double or triple mutations on the protein stability and to determine the ΔΔG values for each protein. For mutations in the Spike protein binding site to RBD from ACE2, the difference in protein stability between Spike protein:RBD complex and Spike protein alone is indicated. For in silico docking, the HawkDock server (http://cadd.zju.edu.cn/hawkdock/) was used to conduct docking simulations between our modeled Spike S protein structures and ACE2. All molecular structures were visualized with PyMol.

## Results

We performed real time RT-PCR tests on 32233 individuals, starting on 16.02.2020 until 22.06.2020, and summarized the data in Table 1. We compared our data to public information on national testing relevant for this pandemic, valid at the time of writing.

**Table 1.** Aggregate of the testing data performed at Cantacuzino Military-Medical National Research-Development Institute (INCDMMC) and at national level [9]

| As of 22.06.2020 | INCDMMC data | National data |
| --- | --- | --- |
| Total number of tests | 32233 | 626330 |
| Confirmed SARS-CoV-2<br>(% from total tested) | 1630 (5.05 %) | 24045 (3.83%) |
| Negative SARS-CoV-2<br>(% from total tested) | 30603 (94.95 %) | 602285 (96.17 %) |

A recent analysis of risk factors in Romanian patients was performed for a subset of 842 deceased individuals or 7.1 % out of a subset of 11790 confirmed cases (data accessed on May 23^rd^, 2020) [10]. For both sexes, patients with pre-existing cardiovascular conditions were 2.89 times more likely to succumb than patients who were not affected by this type of illness. Importantly, the next categories of patients who died were suffering from chronic renal diseases (death more probable by 2.69) and diabetes (by 2.3 times). Given the high expression and circulating levels of hACE2 in the heart and kidneys, these findings support a mechanism whereby Spike glycoprotein may attach to hACE2 in a dose-response manner. We also note that these and other high risk factors (e.g. chronic lung disease or cancer/other immune deficiencies) are not equally distributed between sexes, with men having the highest risks when suffering from cancer or other immune deficiencies (Odds Ratio of 3.25), while women succumbing mostly when suffering with chronic renal diseases (Odds Ratio of 4.3). This finding may reflect possible gender differences in the levels and activity of ACE2, as has been observed in murine models [11].

### Genetic phylogeny

The sequences of Romanian SARS-CoV-2 sequences showed high (∼ 99.95 identity with Wuhan seafood market pneumonia virus (Genbank accession number: NC_045512.2) and >99.6% with sequences from France, Germany, Greece, Italy, Poland, India, USA. Phylogenetic analysis showed that the Romanian sequences belonged to different clusters (Figure 1). The following sequences were introduced in GISAID (Global Initiative on Sharing All Influenza Data) under accession numbers: EPI_ISL_445220, EPI_ISL_445243, EPI_ISL_447054, EPI_ISL_455468, EPI_ISL_455469, EPI_ISL_455470, EPI_ISL_455471, EPI_ISL_455472, EPI_ISL_455473, EPI_ISL_455474, EPI_ISL_455475, EPI_ISL_455476, EPI_ISL_455477, EPI_ISL_455479, EPI_ISL_467779, EPI_ISL_467780, EPI_ISL_467781, EPI_ISL_467778.

**Fig. 1.**
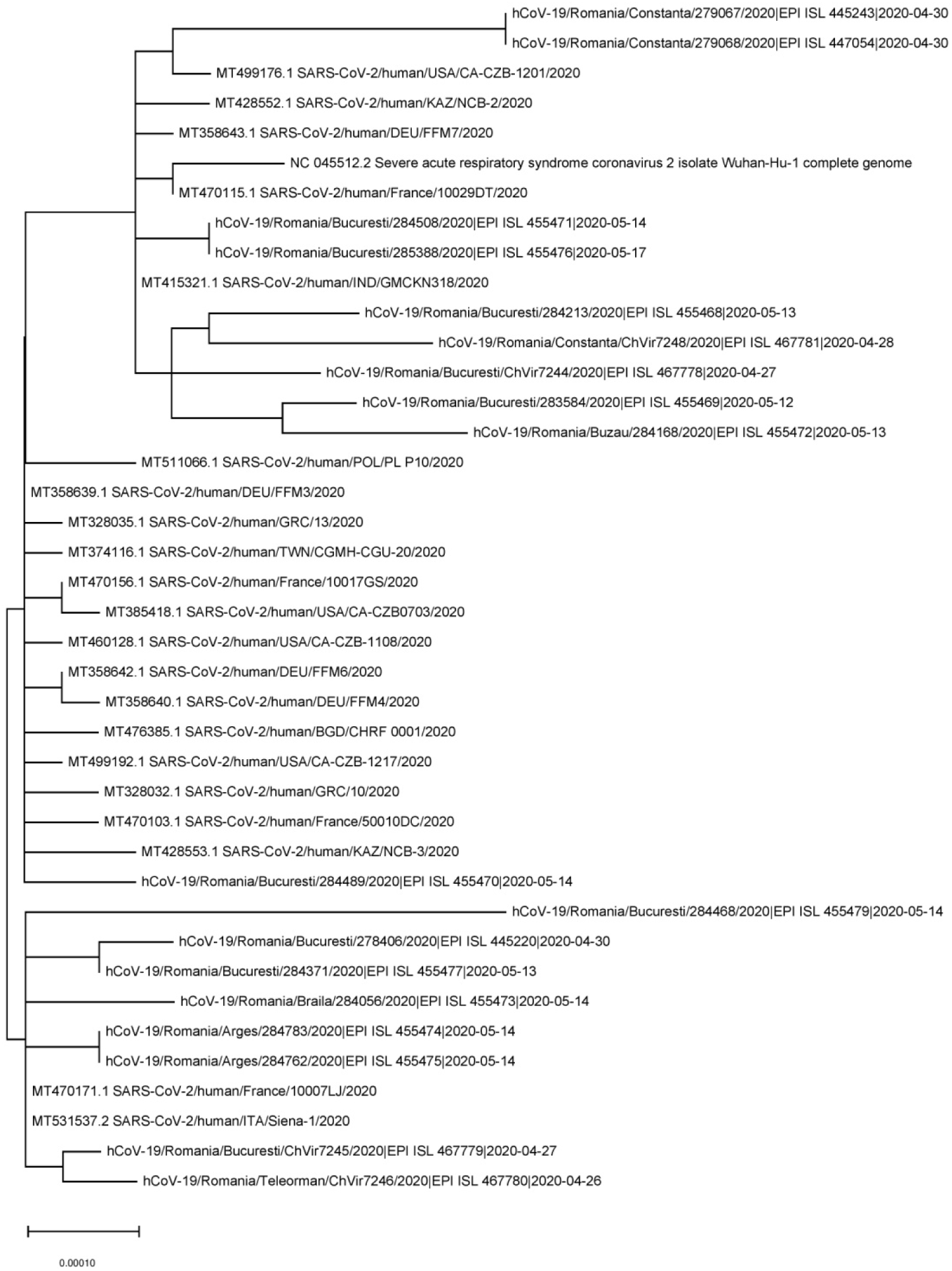
Phylogenetic analysis of 18 SARS-CoV-2 complete genome sequences. The Wuhan reference genome from GenBank accession number: NC_045512.2 and other European/American complete sequences are also shown from different countries (n=21). The tree was built by using the best fitting substitution model (HKY) through MEGA X software.

### Molecular modeling

An overview of the 7 mutations of the Spike proteins detected from Romanian sequence is presented in the structure of Figure 2 and listed in Table 2. We note that only the N439K variant is involved in the binding interface, where the lysine now H-bonds with Gln 325 from the hACE2. Mutants included single, double and triple Spike protein variants. The impact of the mutations on the stability of the complexes made with hACE2 is also indicated in Table 2.

**Table 2.** Key residues involved in the formation of interfaces between ACE2: Spike S variants

| Spike Protein | Viral Interface Residues | RMSD (Å) | TM score | Binding free energy - complex hACE2:variant Spike proteins (kcal/mol) |
| --- | --- | --- | --- | --- |
| PDB 6LZG |  | - | - | **-42.96** |
| Model SARS-CoV-2 RBD [PDB: 6LZG] | K417, G446, Y449, Y453, L455, F456, A475, F486, N487, Y489, F490, **Q493**, G496, Q498, **T500**, A501, G502, Y505 | - | - | - |
| *D614G* variant Spike S overlapped to SARS-CoV-2 RBD | Similar to SARS-CoV-2 RBD | 0.97 | 0.63 | **-9.94** |
| *P521R*, *D614G* variant Spike S overlapped to SARS-CoV-2 RBD | Similar to SARS-CoV-2 RBD | 0.97 | 0.6 | **-27.25** |
| *S98F*, *D614G* variant Spike S overlapped to SARS-CoV-2 RBD | Similar to SARS-CoV-2 RBD | 0.99 | 0.58 | **-26.24** |
| *N439K*, *D614G* variant Spike S overlapped to SARS-CoV-2 RBD | K417, K439, G446, Y449, Y453, L455, F456, A475, F486, N487, Y489, F490, **Q493**, G496, Q498, **T500**, A501, G502, Y505 | 0.96 | 0.61 | **-39.22** |
| *V308L*, *D614G* variant Spike S overlapped to SARS-CoV-2 RBD | Similar to SARS-CoV-2 RBD | 0.97 | 0.51 | **-28.74** |
| *E96D*, *H245Y*, *D614G* variant Spike S overlapped to SARS-CoV-2 RBD | Similar to SARS-CoV-2 RBD | 0.98 | 0.6 | **-67.21** |

**Figure 2:**
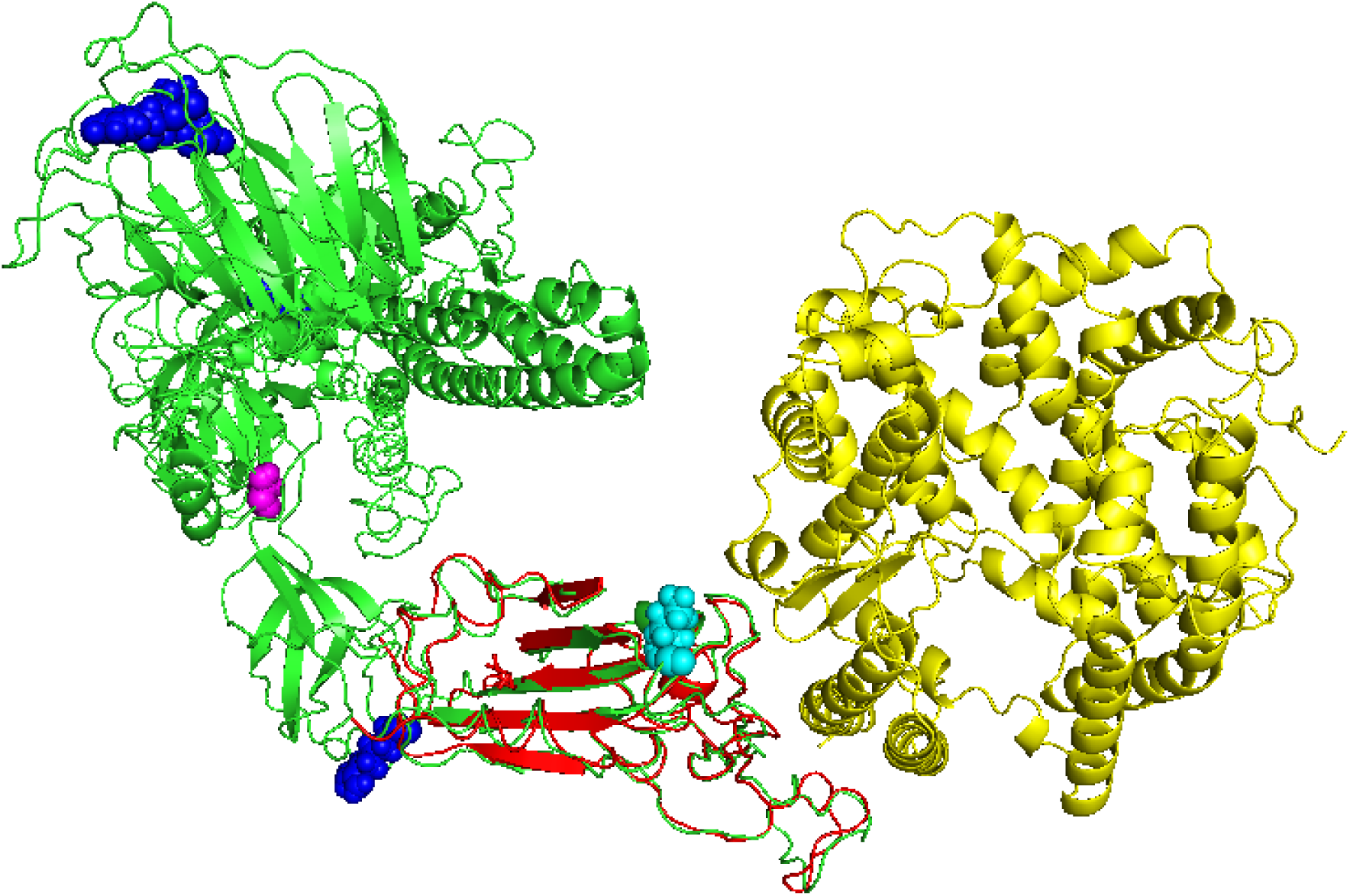
Docking of hACE2 (from PDB 6LZG, in yellow) with models of Spike glycoprotein variants (green) that contains the location of mutations described in this study (blue) with respect to binding partner hACE2. D614G mutation is shown in magenta; N439K mutation, that sits at the binding interface with hACE2 is shown in cyan. The structural overlap between RBD from Spike protein in PDB 6LZG and a constructed model of the mutant Spike protein is shown in red.

We used PDBsum to identify the critical amino acids at the binding interface, and further compiled with PyMol their identity (Table 2), where residues interfacing with multiple H-bonds are shown in bold. TM-scores [0-1] indicating the structural similarity to the overlapped model have been derived with I-TASSER (TM-Scores of 0.5 and above represent a high probability to match similar folds). The impact of the mutations on the stability of the complexes made with hACE2 is also indicated in Table 2. Although all mutants have negative (favorable) binding energy, compared to the native complex, only the triple mutant has a significantly higher value (60% increase). We note that the double mutant *N439K, D614G* that directly affects the binding interface, has only negligible positive effect on binding, but ranks above the rest of the variants.

## Conclusions

Mutations in the spike surface glycoprotein may be conducive to conformational changes, which can translate into changing antigenicity. As such, identifying the amino acids involved in conformational changes of the SARS-CoV-2 spike surface glycoprotein structure is essential to inform on patterns of mutations subject to positive selection. The role of detected mutations on overall thermodynamic stability of the protein variants is thus imperative, as shown in Table 3.

**Table 3.**
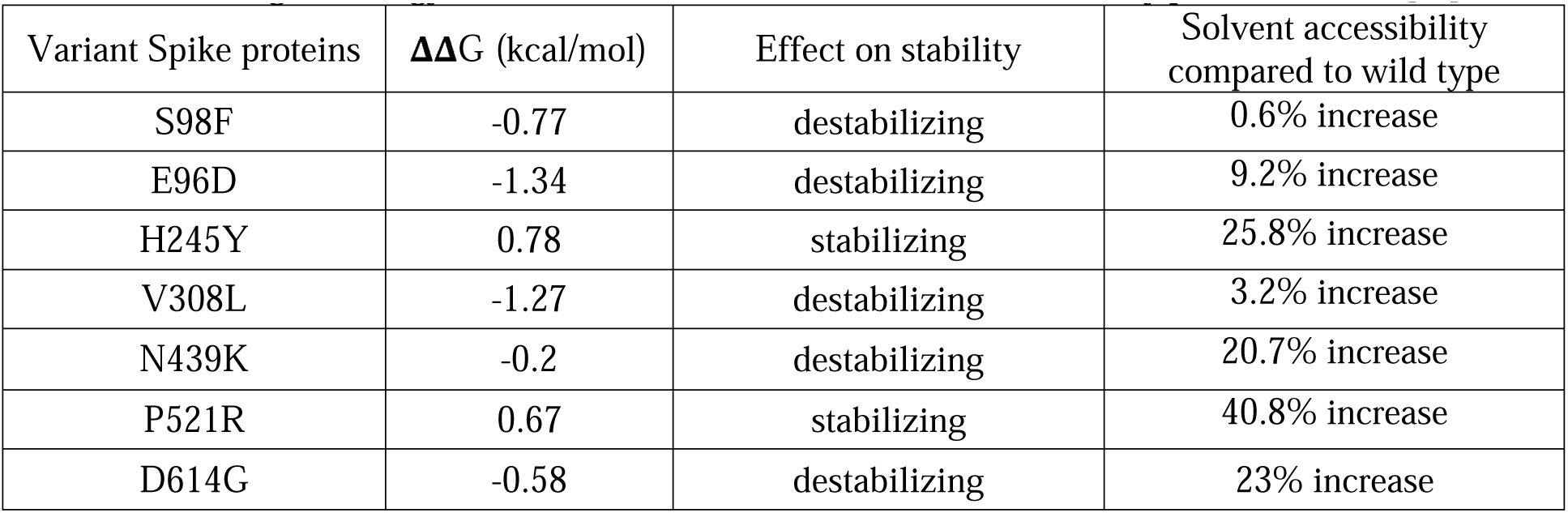
Role of point mutations observed in the genomes on overall protein stability. ΔΔG parameter is a measure of the change in energy between the folded and unfolded states caused by point mutations [12].

| Variant Spike proteins | $\Delta\Delta G$ (kcal/mol) | Effect on stability | Solvent accessibility compared to wild type |
| --- | --- | --- | --- |
| S98F | -0.77 | destabilizing | 0.6% increase |
| E96D | -1.34 | destabilizing | 9.2% increase |
| H245Y | 0.78 | stabilizing | 25.8% increase |
| V308L | -1.27 | destabilizing | 3.2% increase |
| N439K | -0.2 | destabilizing | 20.7% increase |
| P521R | 0.67 | stabilizing | 40.8% increase |
| D614G | -0.58 | destabilizing | 23% increase |

Spike *D614G* variant has been the most widespread mutant encountered thus far in the 2020 genome datasets outside East Europe (3577 counts in GISAID database) [13]. *D614G* mutation is embedded into an immunodominant antibody epitope and is recognized by monoclonal antibodies isolated from recovered individuals, who had been infected with the original SARS-CoV [14]. A very recent experimental study comparing Spike variants *D614G* against *D614S* indicates that the former is less stable thermodynamically, which translates into markedly increased infectivity of ACE-2 expressing cells, consistent with epidemiological data.^15^ Furthermore, pseudoviruses containing both of these variants were neutralized with comparable efficiency by convalescent plasma [14]. However, another recent study demonstrated that Spike variant *D614G* significantly infects ACE-2 expressing cells, when compared to the native Spike protein, and that convalescent sera showed decreased neutralizing activity against *D614G* pseudovirus [16]. Given these factors, this mutant could mediate immune avoidance, although there is as-yet no definite clinical data to correlate the impact of this or any other mutation on infectivity. We note that none of the other mutations we describe have been found thus far in other cohorts, and that other mutations are themselves rare [13]. Tracking mutations in viral Spike glycoprotein continues to be paramount for vaccine and antibody therapy strategies that are currently being developed, and on evaluating new SARS-CoV-2 strains as they emerge and evolve.

## Acknowledgements

We are grateful to Luiza Ustea, Nicoleta Paraschiv and Adrian Cre□u for technical support. We thank the staff working from the Viral Respiratory Infections Laboratory (National Influenza Centre), colleagues from Cantacuzino Military-Medical Research and Development National Institute, the public health staff from the National Centre for Communicable Diseases Surveillance and Control, officials from the Ministry of Health and the Ministry of Defense. Funding support was provided by Rompetrol Group NV (Fundatia pentru SMURD) and the Romanian Association for Shoulder and Elbow Surgery. The funding sources had no role in the study design, analysis or writing of report.

## REFERENCES

[1] Zhou P, Yang XL, Wang XG, et al. A pneumonia outbreak associated with a new coronavirus of probable bat origin. Nature. 2020; 579(7798):270□273.

[2] Zheng F, Tang W, Li H, Huang YX, Xie YL, Zhou ZG. Clinical characteristics of 161 cases of corona virus disease 2019 (COVID-19) in Changsha. Eur Rev Med Pharmacol Sci. 2020; 24(6):3404□3410.

[3] European Center for Disease Control: Rapid Risk Assessment: Coronavirus disease 2019 (COVID-19) in the EU/EEA and the UK– ninth update. Retrieved on June 8^th^, 2020.

[4] Garg S, Kim L, Whitaker M, et al. Hospitalization Rates and Characteristics of Patients Hospitalized with Laboratory-Confirmed Coronavirus Disease 2019 — COVID-NET, 14 States, March 1–30, 2020. MMWR Morb Mortal Wkly Rep. 2020; 69:458–464.

[5] Robson B. COVID-19 Coronavirus spike protein analysis for synthetic vaccines, a peptidomimetic antagonist, and therapeutic drugs, and analysis of a proposed achilles’ heel conserved region to minimize probability of escape mutations and drug resistance. Comput Biol Med. 2020; 121:103749.

[6] Corman VM, Landt O, Kaiser M, et al. Detection of 2019 novel coronavirus (2019-nCoV) by real-time RT-PCR. Eurosurveillance. 2020; 25(3):2000045.

[7] Yang J, Zhang Y. I-TASSER server: new development for protein structure and function predictions. Nucleic Acids Research. 2015; 43: W174–W181.

[8] Williams CJ, et al. MolProbity: More and better reference data for improved all-atom structure validation. Protein Science. 2018; 27: 293–315.

[9] Informare COVID-19, Grupul de Comunicare Strategică, 13 Iunie 2020 (In Romanian). https://www.mai.gov.ro/informare-covid-19-grupul-de-comunicare-strategica-13-iunie-2020-ora-13-00/. Retrieved 14.06.2020.

[10] Popovici O. Risk factors for death in confirmed cases of patients with COVID-19 (In Romanian). https://www.cnscbt.ro/index.php/analiza-cazuri-confirmate-covid19/1791-analiza-epidemiologica-factori-de-risc-deces-cu-covid-19. Retrieved 09.06.2020.

[11] White MC, Fleeman R, Arnold AC. Sex differences in the metabolic effects of the renin-angiotensin system. Biol Sex Differ. 2019; 10, 31.

[12] Kellogg EH, Leaver-Fay A, Baker D. Role of conformational sampling in computing mutation-induced changes in protein structure and stability. Proteins. 2011;79(3):830□838.

[13] Korber B, et al. Spike mutation pipeline reveals the emergence of a more transmissible form of SARS-CoV-2. BioRxiv. 2020.04.29.069054; doi: https://doi.org/10.1101/2020.04.29.069054.

[14] Wang Q, et al. Immunodominant SARS Coronavirus Epitopes in Humans Elicited both Enhancing and Neutralizing Effects on Infection in Non-human Primates. ACS Infectious Diseases. 2016; 2, 361–376.

[15] Zhang L, Jackson CB, Mou H, Ojha1 A, Rangarajan ES, Izard T, Farzan M, Choe H. The D614G mutation in the SARS-CoV-2 spike protein reduces S1 shedding and increases infectivity. BioRxiv. 2020.06.12.148726; doi: https://doi.org/10.1101/2020.06.12.148726.

[16] Hu J, He CL, Gao Q, Zhang GJ, Cao XX, Long QX, Deng HJ, Huang LY, Chen J, Wang K, Tang N, Huang AL. The D614G mutation of SARS-CoV-2 spike protein enhances viral infectivity and decreases neutralization sensitivity to individual convalescent sera. BioRxiv 2020.06.20.161323; doi: https://doi.org/10.1101/2020.06.20.161323.

